# Regulatory mechanism predates the evolution of ant-like swarm intelligence in simulated robots

**DOI:** 10.1101/372391

**Authors:** Ryusuke Fujisawa, Genki Ichinose, Shigeto Dobata

## Abstract

The evolution of complexity is one of the prime features of life on Earth. Although well accepted as the product of adaptation, the dynamics underlying the evolutionary build-up of complex adaptive systems remains poorly resolved. Using simulated robot swarms that exhibit ant-like group foraging with trail pheromones, we show that their swarm intelligence paradoxically involves regulatory behavior that arises in advance. We focused on a “traffic rule” on their foraging trail as a regulatory trait. We allowed the simulated robot swarms to evolve pheromone responsiveness and behaviors simultaneously. In most cases, the traffic rule, initially arising as selectively neutral component behaviors, assisted the group foraging system to bypass a fitness valley caused by overcrowding on the trail. Our study reveals a hitherto underappreciated role of regulatory mechanisms in the origin of swarm intelligence, as well as highlights the importance of embodiment in the study of their evolution.

---

The evolution of complexity is one of the most striking characteristics of life throughout its hierarchy (1, 2). The resulting complexity often entails adaptation to the environment, known as complex adaptive systems (3–5). At the organismal level, theoretical works based on Fisher’s geometric model of adaptation predict that a population of organisms climbing a hill in a fitness landscape evolves a complex polygenic trait based on a few major genes and many minor genes (6–9). Typically, the advent of major genes with large phenotypic effects is followed by the subsequent evolution of minor genes with small phenotypic effects.

Complex adaptive systems beyond the single organismal level, such as multicellularity and social organization, show similar multi-component regulations: The systems seem to take a layered form in which their core components, often considered as evolutionary innovation and hence with large phenotypic effects, are buffered by additional, regulatory components with relatively small phenotypic effects (10). In the study of self-organization in biological systems, such regulation has been referred to as “parameter tuning” (10, 11). Self-organization itself does not guarantee adaptation (10, 12), and previous studies have shown how fine-tuning of endogenous parameters and supplementation of accessory regulation of the core self-organizing systems can lead to variable system outputs (10, 13–16) and make the systems adapt to species-specific (17–19) or within-species changing environments (19, 20). The ability of such adaptive modifications should be attributed to the evolution of regulatory traits that regulate core components. Therefore, for a deeper understanding of the adaptability of systems of biological organization as complex adaptive systems, it is necessary to decipher the multi-trait evolutionary dynamics leading to extant systems (21).

In this study, we address this issue by considering social insect colonies as a model system. Social insects stand at one of the pinnacles of biological complexity. Their colonies are characterized by highly coordinated systems, often likened to swarm intelligence, in which microscopic interactions of nestmate individuals collectively produce diverse macroscopic phenomena (10, 22–24). A typical example is found in mass foraging by ants with trail pheromones (25, 26). The core component of this system is indirect pheromone communication among nestmates according to the following algorithm: Once a worker finds food, she puts a chemical marker on the ground while carrying the food to her nest; nestmate workers are recruited to the marker and follow the trail toward the food, and lay the same marker to reinforce the trail. The system is strengthened by the balance between positive (trail reinforcement) and negative (trail decay) feedbacks depending on changing food availability. The algorithm of worker’s behavior has been applied to solve computational problems such as the traveling salesman problem, known as Ant Colony Optimization (27).

We took a constructive approach with both robotic and computational systems that mimic this foraging system, focusing on its logistic aspects (28, 29). These artificial systems can provide not only controlled experiments that would be impossible in real organisms, but also greater realism through embodiment, which is often abstracted in analytical and purely simulational studies (30–33). Such contributions include the demonstration of evolutionary emergence of cooperative behaviors (34–38) and rigorous testing of kin selection theory (39). During the development of the real robotic system, we faced a problem of how to deal with overcrowding on the trail. Because the use of the trail inevitably puts robots (as it does ants) into traffic-jam-like overcrowding, an accessory regulation that supports efficient pheromone communication is required in both systems. As a solution to the robotic overcrowding, we heuristically introduced a set of collision-processing behaviors in the robots (29, 40). These behaviors constitute an overall “traffic rule”, such that inbound (food-to-nest) robots are always given priority over outbound (nest-to-food) robots. Interestingly, similar collision-processing behavioral rules have been reported in some ants (41; see Discussion). Although the ants and our robots are obviously different in many respects, the two systems share the same property of the layered complexity: the core component (pheromone-mediated group foraging) is supported by regulatory traits (traffic rule). Therefore, we asked how such complex adaptive systems, supplemented with accessory regulations, were achieved through adaptive evolution. In accordance with the conventional notions of polygenic traits, it seemed reasonable to assume that such regulations evolve after the system’s core component has been established, simply because the former rely on the latter for their functions.

Using a simulated system that precisely modeled the dynamic properties of real robots (29), we first confirmed that the algorithm of pheromone communication alone was not sufficient to establish effective recruitment due to the overcrowding on the pheromone trail. The additional traffic rule was required to solve this problem by keeping the robots from being stuck on the trail. Next, we performed evolutionary population genetic simulations that allowed for mutation and selection to occur simultaneously in pheromone responsiveness and collision-processing behaviors of the robots. In most simulation runs, the collision-processing behaviors did not arise as a consequence of pheromone communication, but arose in advance, taking a form of selectively neutral behaviors in the absence of swarm intelligence. The behaviors then assisted the pheromone responsiveness trait to arise and become fixed in the population. Finally, we confirmed the above results with a population genetic analysis hybridized with simulated distributions of fitness values.

## Results

Our robots searched for the location of “food” in a rectangular field (900 mm × 9000 mm) surrounded by walls and enclosing their “nest-site” (Fig. 1a). The field setup was originally intended to facilitate observation of robots on the trail, but it might also be applied to the traffic flow inside ant nests, where numerous ants have to manage overcrowding along their underground galleries. The algorithm for foraging behaviors (29) is described as state transitions among three behaviors: *S*_1_, searching; *S*_2_, carrying food (inbound) and recruiting (laying scent); and *S*_3_, being recruited (outbound, following scent). State *S*_3_ is functional only in the presence of the ability to detect pheromones (Fig. 1b). As an accessory regulation, we implemented a collision-processing behavior: When two robots collide, both take one of two reactions: “Stay” (stop moving for a given time) or “Leave” (move backward by a given distance). We assumed that the robots’ reactions depend on their state (for different assumptions, see Discussion). When colliding robots with different states take different reactions, they can be regarded as obeying a traffic rule, i.e., the robot with the reaction “Leave” gives priority to the robot with the reaction “Stay” (Fig. 1c). To allow for the robotic swarms to evolve, we made a simple assumption that each robot has its own haploid genome consisting of four loci, *b_1_, b_2_, b_3_,* and *p*, each of which has a binary allelic state (0 or 1). The resulting multilocus genotype is described as {*b_1_,b_2_,b_3_;p*}. The locus *b_i_* (*i* ∈ {1, 2, 3}) defines the collision-processing behavior (i.e., Stay = 0, Leave = 1) taken by the robot with state *S_i_*. That is, if a robot is currently in state *S_i_*, then its collision behavior is *b_i_*. The locus *p* defines the ability to detect pheromone.

**Figure 1.**
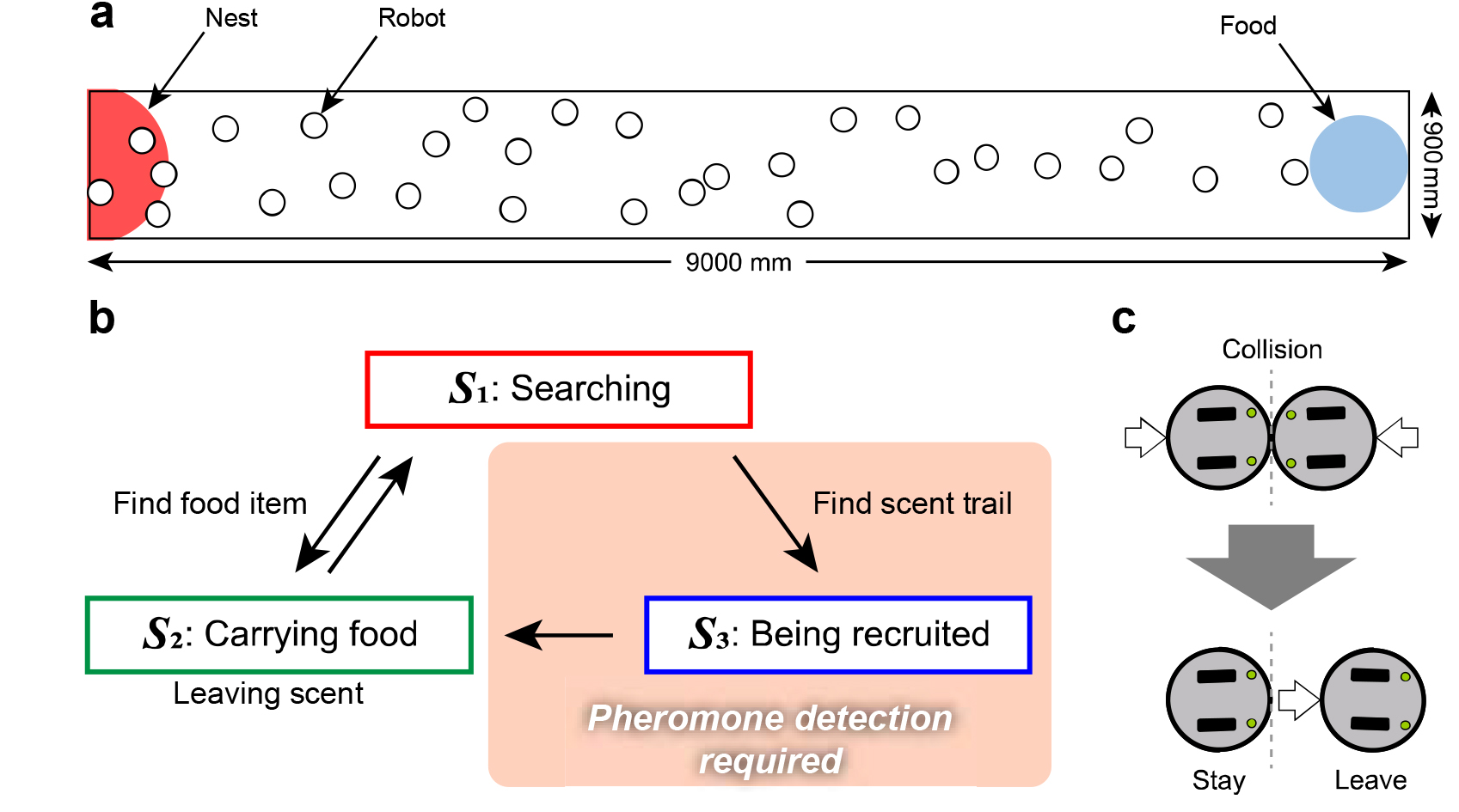
Experimental setup. (**a**) The foraging arena used in the simulations. (**b**) The behavioral algorithm for individual robots with their state transitions (*S*_1_–*S*_3_). (**c**) The collision-processing behaviors.

We measured biological fitness of simulated clonal swarms resulting from their multilocus genotypes ({0,0,0;0} – {1,1,1;1}, 2^4^ = 16 in total) to map a multidimensional fitness landscape. The total number of times the robots go back to their nest from the food was considered as a measure of swarm fitness. The ability to detect pheromone (*p* = 1) alone did not result in higher fitness, although recruitment (*S*_3_) occurred (illustrated as the presence of orange bars in Fig. 2). The reaction “Leave” at locus *b*_1_ contributed to the fitness increase regardless of the presence of pheromone responsiveness. Interestingly, particular sets of behavioral genotypes together with the pheromone responsiveness remarkably improved the swarm fitness, which was achieved through successful recruitment. Among the 16 multilocus genotypes, the genotype {1,0,1;1} showed the highest swarm fitness, in which a traffic rule was established with outbound robots (*b*_3_ = 1) giving priority to inbound robots (*b_2_* = 0) on the pheromone trail. This result was consistent with our previous real robot experiment (40). To confirm that the traffic rule helped the swarms to avoid overcrowding, we counted the number of collisions during each simulation run. The traffic rule on the pheromone trail (*b_2_* = 0, *b_3_* = 1), together with *b_1_* = 1, strengthened the pheromone communication by reducing the occurrence of collision (Supp. Fig. 1, red arrow).

**Figure 2.**
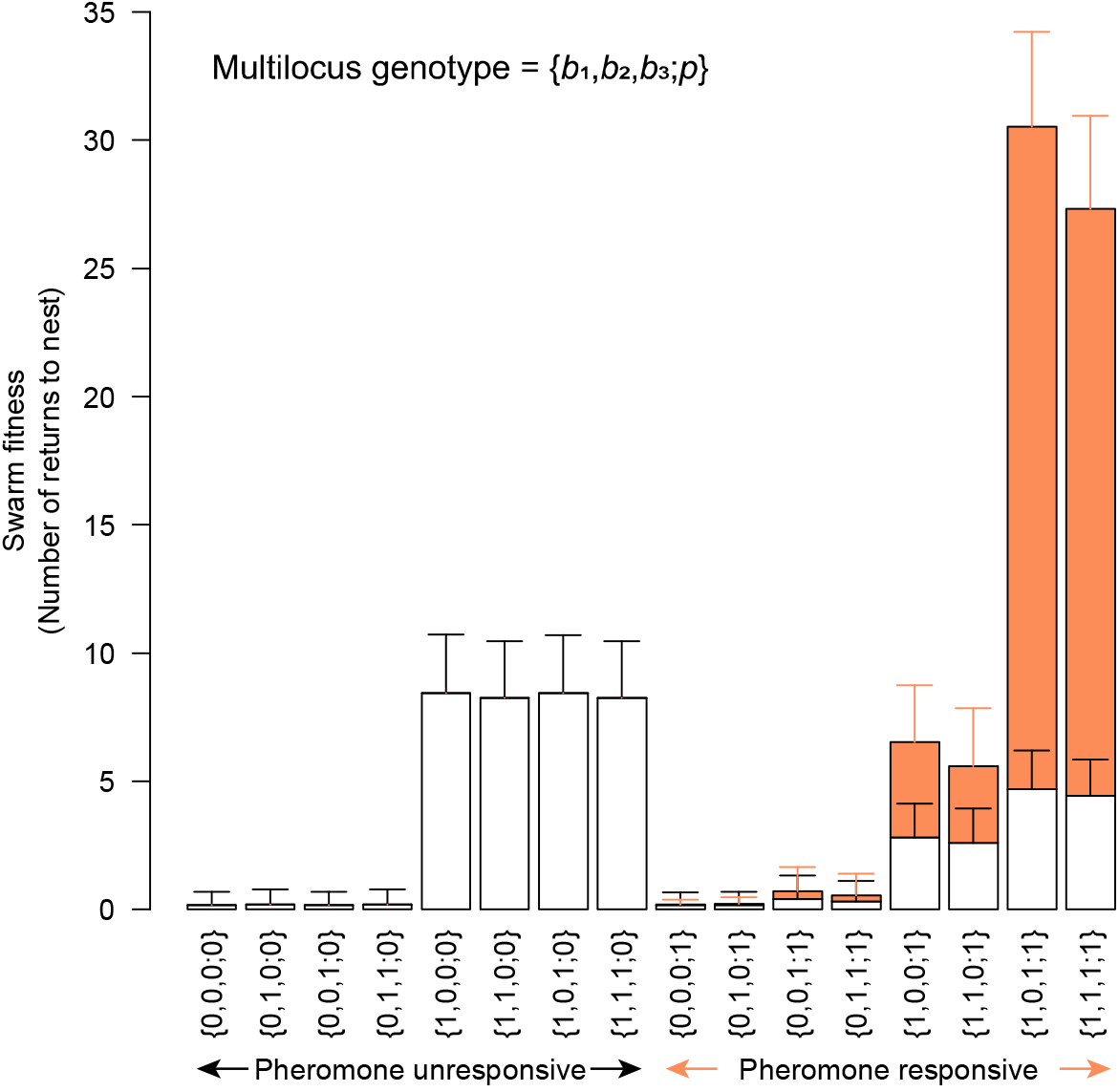
Measures of group foraging performance. Total number of foraging bouts per swarm as a measure of swarm fitness. Bars indicate that the food was found by robots in state *S_1_* (white, searching independently) or *S_3_* (orange, being recruited). Error bars show SD (*n* = 100 000 trials each).

We next conducted population genetic simulations to trace an evolutionary trajectory of the robotic population that initially had the genotype {0,0,0;0}, that is, without pheromone communication and priority-giving behaviors. For simplicity, all robots in a single swarm were assumed to have the same genotype, and genetic variation was permitted only among swarms. In terms of sociobiology, only a single robot in the swarm clonally (with mutations) produces foundress robots of the next generation depending on the swarm fitness, each of which then clonally reproduces (without mutations) to form a new swarm. Consequently, the average relatedness of nestmates within a swarm is strictly 1.

Throughout the simulation replicates, as expected, the populations eventually became fixed at the genotype {1,0,1;1}; that is, the swarms successfully evolved the pheromone-mediated swarm intelligence together with the traffic rule (Supp. Movie 1). Then we inspected the detail of each evolutionary trajectory. The population of swarms first became dominated quickly (frequency ≥ 0.985, fixed in most cases, Supp. Table 1) by the genotype {1,0,0;0}, in which the reaction “Leave” by randomly searching robots (*b_1_* = 1) was not related to pheromone communication (Supp. Movie 1). Starting from the population dominated by the genotype {1,0,0;0} and assuming the shortest path, the robot swarms had to experience 0→1 mutations at both loci *b_3_* and *p* before fixation of the genotype {1,0,1;1}. Surprisingly, in most (44/50) of the simulated genealogies leading to the final genotype {1,0,1;1}, the 0→1 mutations occurred first at the behavior locus *b_3_*, followed by the pheromone-responsiveness locus *p* (Fig. 3a). The intermediate genotype {1,0,1;0} never became fixed; it served as the source of the final genotype when it had an intermediate (0.005– 0.685) frequency in the population (Fig. 3b, Supp. Table 1). Among the rest of the simulated genealogies, five had the 0→1 mutation occurring first at locus *p* and then at locus *b_3_*.

**Figure 3.**
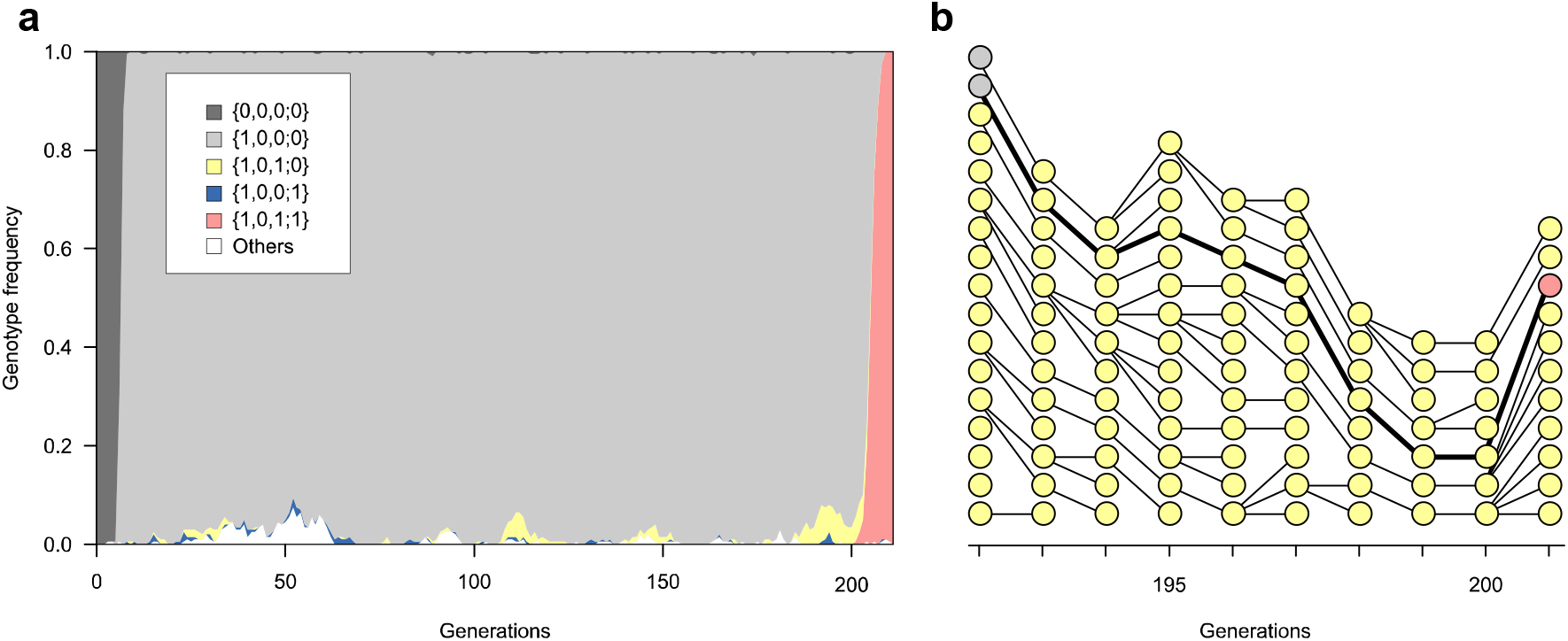
Simulated evolutionary dynamics. (**a**) In most cases, the population of the ancestral genotype {0,0,0;0} (dark gray) was quickly taken over by the genotype {1,0,0;0} (light gray); the genotype {1,0,1;0} (yellow) as well as other genotypes then arose but remained at low frequencies; and the final genotype {1,0,1;1} (red) arose from {1,0,1;0} and quickly became fixed. The whole population showed a state transition from all-{1,0,0;0} to all-{1,0,1;1}. (**b**) Selected genealogies of the genotype {1,0,1;0} during generations 192 and 201 shown in a. Circles denote swarms, with colors as in **a**. This includes the genealogy leading to the final genotype {1,0,1;1}. The regulatory trait *b_3_* = 1 predated the evolution of pheromone responsiveness trait *p* = 1 along the focal genealogy.

The intermediate genotype {1,0,0;1} remained at a low frequency (0.01–0.02). One genealogy did not take the shortest path from {1,0,0;0} to{1,0,1;1}; its sequence was {1,0,0;0}→ {1,1,0;0}→{1,1,1;0}→{1,1,1;1}→{1,0,1;1}. Mutations at the behavior loci were observed both before and after the 0→1 mutation at locus *p*. The irregularity can be explained by an additional 0→1 mutation at locus *b_2_* before the 0→1 mutation at locus *b_3_* while the dominant pattern of evolutionary precedence of *b_3_* = 1 over *p* = 1 remained intact.

The observed bias toward evolutionary precedence of the regulatory trait over the core pheromone-responsiveness trait can be explained using the fitness landscape as follows (Fig. 4a): One of the two possible intermediate genotypes, {1,0,0;1}, actually had lower swarm fitness than the genotype {1,0,0;0} because of overcrowding on the pheromone trail (Supp. Movie 1), whereas the other intermediate {1,0,1;0} gave the same fitness in principle because locus *b_3_* gives a selectively neutral trait by definition, as long as the pheromone-responsiveness is absent (i.e., *p* = 0) (Supp. Movie 1). Consequently, the genotype {1,0,1;0} is more likely to arise first.

**Figure 4.**
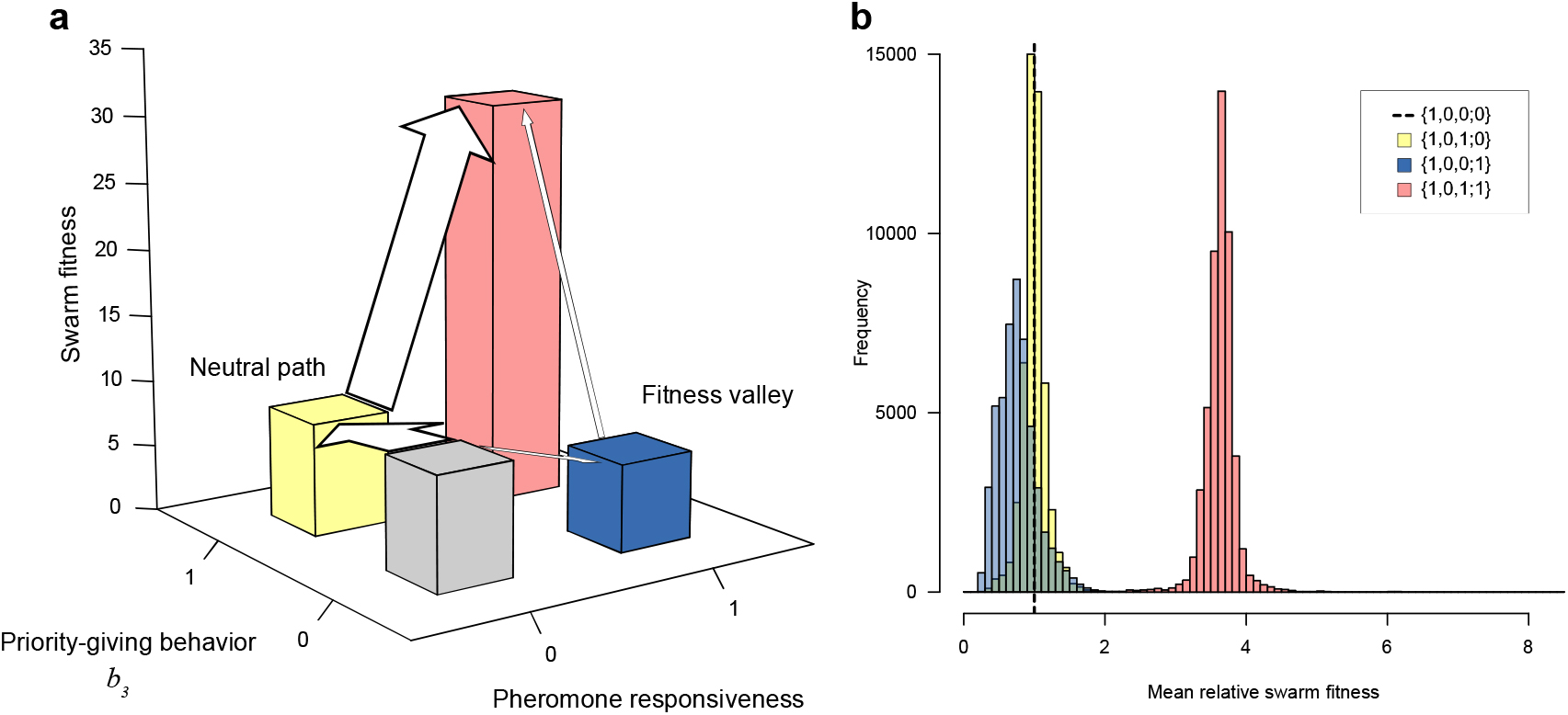
(**a**) Schematic diagram of fitness landscape involving the core (*p*) and regulatory (*b_3_*) traits of the swarm intelligence. Starting from the genotype {1,0,0;0} (light gray), the swarm fitness of the genotype {1,0,0;1} (blue) is generally lower than that of the genotype {1,0,1;0} (yellow), making the latter more likely to arise first (depicted by the thicker arrow) and to serve as the precursor of the final genotype {1,0,1;1} (red). Height of the bars corresponds to swarm fitness in **Fig. 2.** (**b**) Resampled distribution of relative swarm fitness of the neutral intermediate {1,0,1;0} (yellow), inferior intermediate {1,0,0;1} (blue), and final {1,0,1;0} (red) genotypes, compared with the original genotype {1,0,0;0} (- - -). The distributions were incorporated into the mathematical analysis.

The evolutionary process with selectively neutral or even inferior intermediates is known as stochastic tunneling (42, see Discussion). To evaluate how common the observed evolutionary precedence of the regulatory trait (45/50; note that we tentatively included the irregular genealogy described above) is, we applied the population genetic formulation of stochastic tunneling (43) that analytically yields the point estimate of waiting time to fixation of the genotype {1,0,1;1}. Starting from the population fixed at the genotype {1,0,0;0}, the evolving population takes one of the two possible shortest evolutionary paths—that is, the path with the intermediate genotype {1,0,1;0} (*b_3_* = 1 first) and that with the alternative intermediate {1,0,0;1} (*p* = 1 first)—and the realized path could be predicted as the one with the shorter waiting time by comparing respective estimates. Among the parameters of the analytical model were the relative fitness (compared to the original genotype {1,0,0;0}) of the neutral intermediate {1,0,1;0}, the inferior intermediate {1,0,0;1}, and the final genotype {1,0,1;1}, which had to be derived empirically from the evolutionary simulations. Therefore, we incorporated resampled distributions of relative mean swarm fitness (Fig. 4b) into the analytical model (see Methods and Supp. Note for details).

We generated 1000 sets of 50 evolutionary outcomes that the populations were expected to show. The 50 outcomes were typically dominated by those with the intermediate genotype {1,0,1;0}, and their frequency distribution (calculated over the 1000 sets) included the observed frequency (45/50) at the borderline of the 95% interval between the 2.5th (34) and 97.5th (45) percentiles (Supp. Fig. 2). The rare outcomes with the inferior intermediate (i.e., the precedence of pheromone responsiveness) were explained by the stochasticity in the swarm fitness that sometimes resulted in the higher mean relative swarm fitness of this genotype than of the neutral intermediate (Fig. 4b). Moreover, the waiting time to fixation averaged over the 50 runs (157.68 generations) again fell within the 95% interval between the 2.5th (145.54) and 97.5th (354.76) percentiles of the distribution of the mean waiting time (Supp. Fig. 3). These analyses support the view that the phenomenon observed in our evolutionary simulations is well captured quantitatively by the theory of stochastic tunneling, and that the evolution of swarm intelligence in our robotic system was facilitated by this evolutionary process.

## Discussion

One of the long-standing debates in evolutionary biology ever since Charles Darwin is based on the observation that the complexity of life we can find today seems too sophisticated to be achieved through gradual evolution (44, 45). Advocates of saltationism or macromutationism claim that gradualism cannot account for the incipient stages of complex adaptive traits because the intermediate forms must be maladaptive (44). The debate has consequently motivated further work on the genetic and developmental basis of organismal complexity, leading to the rise of present-day evolutionary developmental biology (reviewed in e.g., Refs. 46, 47). The current consensus is that both views are correct depending on systems, and that the real evolutionary processes are fairly complex (48). These contrasting views on the origin of complexity can also be applied to the evolution of social systems, which motivated our present study. In our robotic swarms, the intermediate maladaptive stage corresponds to the pheromone-based recruitment without the traffic rule. By taking a standard population genetic approach, we demonstrated that the resulting fitness valley can be dynamically bypassed through a selectively neutral alternative, that is, cryptic priority-giving behavior without pheromone-based recruitment. Here, the regulatory mechanism was not an evolutionary follower of the core component; instead, it preceded the establishment of the core component and assisted it. The evolutionary precedence of some regulatory mechanisms originated as selectively neutral traits (neutral from their regulatory functions) might be a general component feature of complex adaptive systems, as long as the systems’ core components alone are maladaptive and the accessory regulations rely for their functions on the core components. It does not rule out the possibility that the regulatory mechanisms can arise after the establishment of the core components. The resulting layers of accessory regulations should contribute to the complexity of life.

### Exaptive origins of regulatory mechanisms

The role of the cryptic regulatory mechanisms can be understood by employing the structuralist concept of exaptation (49), a formal definition of so-called preadaptation. Exaptation refers to the evolution of traits that had originated by the selective advantage other than their current use (pre-aptation) or that had originally been non-adaptive (non-aptation or spandrel); both were subsequently co-opted into current adaptive use. The regulatory behavior of our system may provide an example of non-aptation because, by definition, the trait *b_3_* = 1 was selectively neutral at its origin. It should be noted here that we can also consider different genetic coding of the three behavioral traits (*b_1_*–*b_3_*). On the basis of the biologically reasonable assumption that the trait *b_3_* = 1 (or *b_2_* = *b_3_* = 1) is a pleiotropic byproduct of *b_1_* = 1, again we can expect exaptive precedence (i.e., spandrel or a neutral byproduct of previous adaptation in other contexts, 50) of the regulatory mechanism in this foraging system, otherwise the fitness valley would arise owing to overcrowding (Supp. Note).

In our simulations, the bypassing of the fitness valley was driven by the evolutionary process called stochastic tunneling. This was originally proposed as a mathematical description of cancer initiation (42, 51), and the same mechanism was independently found in an analysis of the evolution of *cis*-regulatory elements (52). Taking cancer initiation as an example, its somatic evolution is characterized by a two-step process leading to biallelic loss-of-function mutations at the tumor suppressor gene. The first mutation at a single allele is either selectively neutral or disadvantageous (through chromosomal instability) for cell proliferation, and the second mutation at the counterpart allele triggers increased proliferation as tumor cells. Through stochastic tunneling, the population of cells can shift from the all-intact state to the all-tumor state without experiencing the all-intermediate state. The situation is very similar to our model, except that the genes involved are more than one in our clonal robots. The population of robot swarms favored the neutral intermediate over the disadvantageous alternative, which was easily explained by the comparison of waiting time estimates between the two tunneling routes. The role of stochastic tunneling in the origins of more complex systems, especially those with recombination (21), epistasis (53–55), or indirect genetic effects (54, 55), deserves further study.

A potential concern about applying our rather retrospective approach (i.e., the genetic encoding was made after the best set of behaviors thus far had been known heuristically, see Introduction) to real complex adaptive systems would be that biological systems cannot tell *a priori* what should serve as regulatory mechanisms before the emergence of a core system. Nevertheless, we can predict that complex adaptive systems should be found more frequently in systems allowing more capacity for neutral genetic variations as a source of exaptation. The role of cryptic genetic variations in the emergence of evolutionary novelty is well acknowledged in current evolutionary biology (56). Such cryptic variations would help a primitive system to avoid crossing the fitness valley by providing selectively neutral alternatives. In the context of social organizations, a previous theoretical study predicted that genes with social effects should harbor more variations within a population owing to weaker selection pressure on indirect genetic effects (57). Meanwhile, computational studies that focused on the architecture of biological systems, such as genetic and neural networks (58–60) and secondary structures of RNA molecules (61), have acknowledged the importance of neutral variations as evolutionary capacitors. By highlighting the importance of exaptation and neutral genetic variations for complex adaptive systems, our study bridges an apparent gap between computational and macro-biological studies on the evolution of biological self-organization. Phylogenetic comparative methods might help to test our prediction empirically by reconstructing multi-trait evolutionary processes that lead to extant complex adaptive systems.

### Convergence of traffic rules between ants and robots

Our simulated robots favored the traffic rule under which inbound robots had priority over outbound robots on the trail. In real social insects, previous studies reported traffic-rule-like behaviors shown by ant foragers along their foraging trails (reviewed in Ref. 41). Examples include three-lane bidirectional traffic flow (62), alternating clusters of inbound and outbound ants facing a traffic bottleneck (63), inbound leaf-laden ants followed by unladen ants (64), and alternative route selection through inbound– outbound behavioral interactions at the junction (65). In those examples, macroscopic traffic flow emerges from microscopic behavioral rules where inbound ants are given priority over less-loaded outbound ants. Using embodied robots supplied with virtual food, our study demonstrated that such traffic rules do have an adaptive significance for efficient logistics, in addition to their mechanism (proximate cause) in which real inbound ants have less maneuverability due to food load (41).

To obey the traffic rule, the trail-following robots (i.e., with state *S*_3_) need to put the priority-giving behavior temporarily above the core pheromone responsiveness. This temporal irregularity becomes evident when the reaction “Leave” often moves the robot away from the pheromone trail (Supp. Movie 1). The priority-giving behavior is released after the direct experience of collision with inbound robots, while the pheromone responsiveness can be regarded as following socially available information of food location. Therefore, the adaptive use of the traffic rule might be a situation whereby the robots flexibly prioritize direct social information (collision) over indirect social information (pheromone trail) depending on their internal state during behavioral decision-making. Recent studies have revealed how ants make decisions under such conflicting information (reviewed in Ref. 66). The use of multiple information sources and their integration during collective decision-making would be of particular interest in future study.

An advantage of taking constructive approaches with embodied agents is that we can incorporate physical consequences derived from properties of the agents and their abiotic and biotic interactions. Some of them might manifest as physical constraints hindering adaptation such as the overcrowding we observed, but a more positive aspect would be greater, often unpredictable, degrees of freedom (or evolvability) supplied to the dynamic systems. Previous studies of experimental evolution with swarm robots have revealed various aspects of such consequences, including coordinated behaviors (e.g., 67) and self-organized division of labor (68) (see also Introduction). The evolutionary convergence of traffic rules between ants and our robots, together with those earlier studies, clearly indicates that a collaboration between macro-biology and swarm robotics provides a promising avenue to elucidate the evolutionary and developmental processes leading to the complexity of social life, as well as a hopeful engineering application to solve our real-world problems.

## Methods

The basic algorithm for the pheromone-mediated group foraging behavior (29) has been well validated by comparison of the dynamic properties between simulated and real robot systems (28, 29, 40). In brief, once a searching robot (state *S_1_*) finds food, it starts to secrete a chemical compound on the ground while returning to its nest with a virtual food item (state *S_2_*). If the robots can detect the resulting chemical trail as a trail pheromone, they then follow the trail toward the food (state *S_3_*, regarded as successful recruitment) (Fig. 1b). The priority-giving behavior is described in the main text. The real robot system, named ARGOS-02 (“Ant”elligent Robot Group Operating System, note that our system is different from another swarm robot system named ARGoS, 69), used an aqueous solution of ethanol as the trail pheromone. ARGOS-02 is a modified version of the original ARGOS-01 (28), using two sets (instead of the original one) of micro-pumps and tanks to secrete the pheromone at arbitrary concentrations.

### Experimental setup

A simulated swarm consisted of *n* = 30 robots, which were provided with a rectangular field of 900 mm × 9000 mm surrounded by walls, together with a food source (ø 300 mm) and a nest (ø 1000 mm) on opposite ends (Fig. 1a). The food emitted light, which a robot could detect within a radius of 600 mm from its center. The nest location was made available to all robots by provision of an infrared light directly above it. The initial positions of the robots were set randomly on the field, and each swarm was allowed to forage for 12 000 time steps (20 min for the real robots).

### Evolutionary simulations

The evolving population consisted of *N* = 200 non-interacting swarms (i.e., no resource competition among swarms). We assumed a Wright–Fisher population with a constant size. The genetic coding of the traits is described in the main text. As a prior state to the pheromone-related trait *p*, we implicitly assumed that the inbound robots had already secreted some chemical substance on the ground, and that pheromone communication was achieved upon acquisition of the detection ability as a cue (i.e., 0→1 mutation at *p*). This assumption is based on the “sender–precursor model” of signal evolution (70). A swarm was selected as a clonal parent for the next generation’s swarm in proportion to its fitness, so that a swarm with the better foraging performance had the greater chance of being selected as a parent swarm. In the next generation, *b_i_* and *p* of all genomes in an offspring swarm mutated to the other value (0→1 or 1→0) with a probability μ = 0.001. The low value of mutation rate, compared to conventional evolutionary simulation studies used in computer sciences, was intended to approximate to real biological systems (i.e., it should take relatively long generations for a polygenic system to reach the optimal state). On the basis of the preliminary observations that the genotype {1,0,1;1} is uninvadable by the other genotypes, each simulation run continued until the population became fixed at that genotype. Genotypes of the parent swarms of the genotype {1,0,1;1} that became fixed were determined by direct assessment of the genealogies.

### Analytical solutions of time to fixation

We considered the time until the genotype became fixed {1,0,1;1}, starting from the population of the genotype {1,0,0;0}. The time to fixation can be obtained analytically by using population genetic formulations (43). Four kinds of genotypes were considered: the original genotype {1,0,0;0}, the two intermediates {1,0,1;0} (neutral) and {1,0,0;1} (inferior), and the final genotype {1,0,1;1}.

We calculated the time (generations) until the final genotype became fixed, starting from the population with the original genotype. The original-to-intermediate transition and the intermediate-to-final transition at a swarm occur with the same probability μ, which corresponds to the mutation rate of a single locus. We obtained distributions of the relative fitness (compared to the original) of the neutral intermediate (denoted by *r_0_*), the inferior intermediate (*r*–), and the final genotype (*a*) using data from our evolutionary simulations, as described in Supp. Note.

The probability of tunneling can be given by *T* = *N*μ[1 – *U*(*r_x_*)](1 – *v*_1_), where *N* is the number of swarms, *U*(*r_x_*) is the probability of fixation of the intermediate genotype with relative swarm fitness *r_x_* (*x* ∈ {0, –}), and *v*_1_ is the probability of non-appearance or extinction of the final genotype lineage arising from a single swarm of the intermediate genotype with relative swarm fitness *r_x_* (43). The expected time until the final genotype is given by:

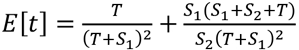
where *S*_1_and *S*_2_describe the probabilities that primary and secondary mutations, respectively, become fixed. In the equation, the first term represents the contribution from tunneling paths (original → final) and the second term the contribution from the sequential paths (original → intermediate → final) (43). Given that a shorter time results in a higher probability of realization, the evolutionary paths are expected to go through the intermediate genotype with the highest relative fitness when there are more than one candidate tunneling paths. See Supp. Note for details.

The main codes of the simulations were implemented in C and R. To shorten the calculation time, we used parallel computation realized by OpenMP. The evolution of swarms was managed by Python programs. For the analytical calculation, we used Mathematica, especially for large matrix calculations.

### Code availability

The source code is available at https://github.com/SWARM-ARGOS/.

## Acknowledgements

We thank H. Shimoji, N. Mizumoto, M. S. Abe, R. Yamaguchi, and T. Tsuchimatsu for discussions. This study was partly supported by Grants-in-Aid for Scientific Research 15K18609 and 17H01249 from the Japan Society for the Promotion of Science (S.D.) and the Hamamatsu Foundation for Science and Technology Promotion (G.I.).

## Author contributions

R.F. developed simulated and real robot systems; S.D. conceived research; G.I. designed evolutionary analyses; G.I and S.D. developed programs for mathematical analyses; R.F., G.I. and S.D performed analyses; S.D wrote the draft, which was edited by R.F., G.I. and S.D.

## Competing interests

The authors declare no competing financial interests.

